# Convolutional neural networks for automated CMR image segmentation in rats with myocardial infarcts

**DOI:** 10.1101/2020.12.01.405969

**Authors:** Andrea Gondova, Magdalena Zurek, Johan Karlsson, Leif Hultin, Tobias Noeske, Edmund Watson

## Abstract

In translational cardiovascular research, delineation of left ventricle (LV) in magnetic resonance images is a crucial step in assessing heart’s function. Performed manually, this task is time-consuming and prone to inter- and intra-reader variability. Here we report first AI-based tool for segmentation of rat cardiovascular MRI. The method is an ensemble of fully convolutional networks and can quantify clinically relevant measures: end-diastolic volume (EDV), end-systolic volume (ESV), and ejection fraction (EF) automatically.

Overall, our method reaches Dice score of 0.93 on the independent test set. The mean absolute difference of segmented volumes between automated and manual segmentation is 22.5μL for EDV, 13.6μL for ESV, and for EF 2.9%. Our work demonstrates the value of AI in development of tools that will significantly reduce time spent on repetitive work and result in increased efficiency of reporting data to project teams.

## Introduction

Cardiac magnetic resonance (CMR) imaging in small animals is one of the most important methods for cardiac function assessment in translational cardiovascular research which increasingly relies on use of tailored rodent models, such as rats and mice, due to the similarities of their cardiac morphology, physiology, function, healing, and post-injury remodeling with humans. CMRI’s non-invasive nature and ability to differentiate tissue types with high spatial and temporal resolution make this method especially suited for a longitudinal follow up of the progression of the cardiovascular disease or response to treatment during the preclinical drug development without the need to sacrifice animals at intermediate points in the study [1,2].

Typically, defects in rodent cardiac structure and function are manifested as changes in left ventricular volume and decrease in ejection fraction. Functional assessment of the left ventricle (LV) is thus the most common task addressed by preclinical teams and currently requires manual delineation of the ventricle, at minimum, at end-diastole (ED) and end-systole (ES) phases of the cardiac cycle [3].

During the analysis, a cross-sectional view of the rodent heart is captured at equal time intervals during one cardiac cycle and ED and ES phases are determined from the recording upon visual examination. Human readers then proceed by tracing LV contours manually in a slice-by-slice manner. The reconstructed ventricular ED and ES volumes are used to calculate LV volumes and ejection fraction.

Segmentation, when performed manually by an experienced reader, requires on average 20 minutes per phase. As typical preclinical study can include up to 50 animals which are followed at multiple timepoints, the task can add up to hours of analysis time lost on tedious, repetitive work. Moreover, although human readers are usually very consistent with low intra-reader variability, the delineation of LV contours are significantly influenced by readers’ experience and knowledge and results in high inter-reader variability between different human experts [4,5,6]. Thus, automated end-to-end segmentation tool that could standardize and speed up the analysis of rodent CMRI datasets is highly desirable.

Examples from clinical setting demonstrate that even though accurate CMRI segmentation is still acknowledged to be difficult, the quality of the cardiac segmentation tools that rely on semi-automated and fully automated approaches have been steadily improving, reaching close to human performance in accuracy of segmentation of difficult human CMRI datasets. This is in no small part due to the developments of deep learning, convolutional neural networks (CNN) especially, and the surge in their uptake within the computer vision community driven by an improved computational capability of general purpose graphics processor units (GPUs) and increased availability of large labeled training datasets [7,8].

CNN-based methods often outperform non-learning based algorithms and traditional machine learning methods which are often criticized for their limited representational ability for the variation and appearance of anatomical organs and their requirement for significant feature engineering and extensive domain expertise [9,10]. In contrast, CNNs’ success has been attributed to their ability to learn hierarchical representations of the raw input data and extract a set of discrimination features automatically. As the raw inputs are passed and processed through the network layers, the level of abstraction of the resulting features increases. Shallow layers grasp local information while deeper layers use filters with broad receptive fields and can capture global information from the image leading to better performance at various imaging tasks [11].

Fully convolutional networks (FCNs) first proposed by Long et al. [12] are a special type of CNNs proposed for image segmentation. FCNs are designed to have a symmetric encoder-decoder structure that can take image of arbitrary size, effectively learn structures in the data and reconstruct the full-sized output that creates a pixel to pixel (or voxel to voxel in case of 3D variants) mask that differentiates between regions belonging to structures of interest vs the background. Given an image, the input is first transformed into a high-level feature representation by the series of convolutional and pooling layers in the encoder path. The decoder than interprets the feature maps and recovers spatial details back to original image dimensions through a series of up-sampling and convolution operations. However, FCN’s simple encoder-decoder structure is somewhat limited in capturing detailed context information necessary for precise segmentation as some features may be eliminated by the pooling layers during the encoding. Because of this, simple FCNs can struggle with reconstructing the segmentation’s finer details and number of different variations on the basic FCN structure were proposed to improve the segmentation accuracy.

The most well-known variant of FCN is the U-net [13]. To boost the segmentation accuracy, U-Net employs skip connections that propagate features from encoder to decoder to help with the recovery of spatial context lost in the down-sampling path. U-net often yields more precise segmentations and several state-of-the-art cardiac image segmentations methods have adopted the U-net or its 3D variants such as 3D U-NET [14] and 3D V-NET [15] as their backbone networks [16,17,18]. We refer the reader to reviews on medical MRI analysis in general [19] and cardiovascular image analyses in particular [10] for more detailed overviews of these applications.

Inspired by results of automated segmentation models in human CMR datasets, we aimed to develop similar approach to efficiently obtain accurate LV segmentations across phases of the cardiac cycle and derive LV volumes and ejection fraction without input of the human in our smaller rodent datasets. To the best of our knowledge, similar end-to-end analysis had not yet been attempted in preclinical cardiovascular setting.

To achieve the desired objective, we leveraged the full spatial information encoded in available volumetric datasets and used networks based on 3D convolutions. There are limitations to this approach as 3D methods drastically reduce the training dataset and have much larger memory requirements compared to their 2D counterparts. However, the loss of context during training on 2D slices rather than whole volumes often results in problems with consistency between neighboring slices, most prominently in apical slices where reported segmentation models sometimes completely fail to detect presence of the ventricle [22, 23]. Theoretically, 3D methods should be more robust to these errors. Our method is inspired by 3D U-net and accepts a full volumetric CMRI data to learn and reconstruct the full-sized voxel-to-voxel mask that differentiates left ventricle from the background. Additionally, we also experimented with a M-net [21]-like architecture which we extended to 3D to leverage the full CMRI volumes.

Like other authors, we improved the quality of final segmentations by using multiple well performing models in an ensemble. As the deployment of large number of models comes with computational costs, we used six best performing models which lead to satisfactory results while maintaining the time-saving benefit of the automated method. Individual, independently trained models used majority voting to assign each voxel to either LV or background. Resulting segmentations were used to calculate volumes of the delineated structure at end diastole and end-systole from which EF was calculated.

As the quality of the segmentation has a direct impact on assessment of the therapeutic agents’ effects in the preclinical study, we performed a thorough validation of the method’s accuracy and reliability. The quality of image segmentations was studied in the independent testing set both visually and using a popular spatial overlap index, Dice score. Additionally, we evaluated the performance in terms of LV ED and ES volumes and ejection fraction. The testing set was also segmented by two human reader which allowed us to assess how well the performance of the automated method compares to inter-reader variability.

## Methods

### Datasets

The training dataset consisted of 636 end-diastolic (ED) and end-systolic (ES) short-axis cardiovascular rat datasets. To achieve heterogenous dataset for model training, the images from sham-operated and rats with subacute and chronic infarcts with LV ejection fraction (EF) range of 67%-41% assessed with Segment V2.2 r6289 (Medviso), were used. All images were acquired using a standard imaging protocol and the same Bruker BioSpin MRI scanner model.

The training dataset consisted of 636 end-diastolic (ED) and end-systolic (ES) short-axis cardiovascular rat datasets. To achieve heterogenous dataset for model training, the images from sham-operated and rats with subacute and chronic infarcts with LV ejection fraction (EF) range of 67%-41% assessed with Segment V2.2 r6289 (Medviso), were used. All images were acquired using a standard imaging protocol and the same Bruker BioSpin MRI scanner model.

### Data pre-processing and augmentation

Size of the available datasets varied in their three dimensions for two reasons. Firstly, during the recoding, the slice thickness is fixed and thus, number of slices can differ between animals depending on the size of the rodent’s heart. Number of slices per dataset within the training set ranged from 11-13. Secondly, readers often crop the areas of interest in the datasets before the manual segmentation which then results in datasets variable in their height and width. As the training requires, at least in every training batch, the images to have the same size, we interpolated the training datasets to the fixed size of 86×98×12 for width, height and number of slices in a stack respectively before training. Next, the image intensities were scaled and centered to that the values lied between 0 and 1. The image processing was perform using scikit-image v0.14.1 package [25].

Additionally, the augmentation of the training dataset was performed using rigid random translations with the maximum shift of 10% of the image height and width. We performed the augmentation progressively and observed that augmenting the dataset 90 times led to improvements of the performance. Further augmentation above this value did not bring additional benefit to the training.

### Training and optimization of the model ensemble

Architecture of our models was inspired by U-net proposed by [13] and it’s 3D variant [14] (Figure 1).

**Fig.1.**
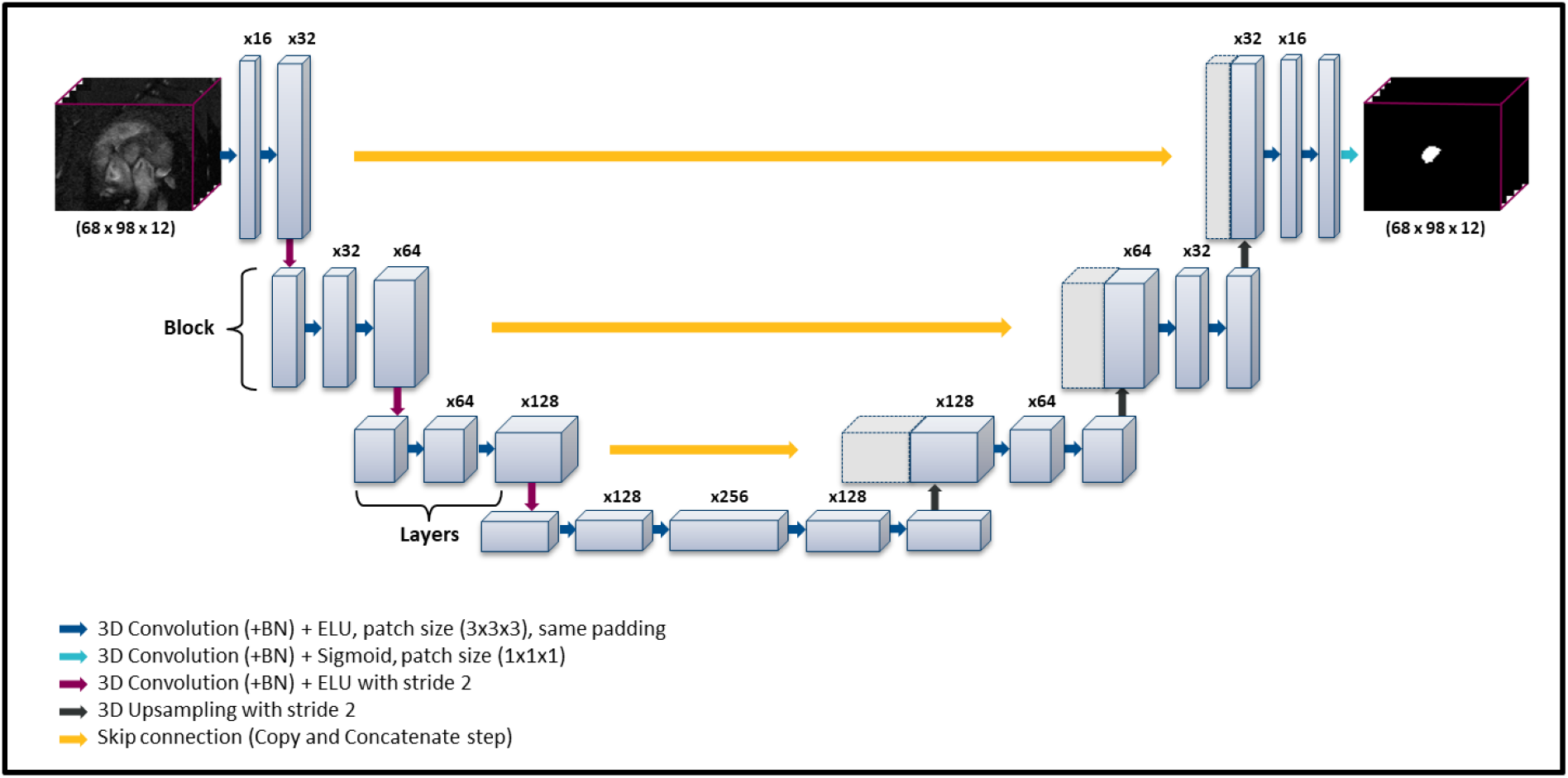
Schematic representation of 3D U-net network architecture. The network accepts a stack of CMRI volumes, learns image features through the encoder, gradually recovers spatial dimensions in the decoder and outputs a predicted mask for the left ventricle (white region) of the same dimensions. Blue boxes represent feature maps with number of maps denoted above the box. Gray boxes represent copied feature maps. For simplicity, we have reduced the number of down-sampling and up-sampling blocks as well as number of layers per block in the diagram.

Generally, the architecture consists of two symmetric encoder and decoder paths that give the network it’s u-shape. The encoder is a typical convolutional network consisting of repeated application of 3D convolutional layers followed by a non-linear activation layer, and a pooling layer that performs the down-sampling of feature maps. Number of feature maps is doubled before each down-sampling. The decoder is symmetrical with the encoder and consists of successive layers where down-sampling step are replaced by up-sampling operators that increase the resolution of their output. The up-sampling is followed by a 2×2×2 convolution that halves the number of feature maps. Additionally, the two arms are interlinked via concatenating skip connections that combine high resolution features from the encoder with the global context in the decoding steps. Thus, local information obtained during down-sampling can be provided to the up-sampling decoding steps and concatenated with the global contextual information in the up-sampling path. This combination of localization and context allows successive convolution layers improve localization and to assemble a more precise output is necessary for good final segmentation. The decoder is followed by a final convolutional layer with sigmoid activation function that outputs a voxel-wise classification of the same resolution as the original input image.

Additionally, inspired by Mehta et al.’ M-Net [21] we extended our U-net by building additional side-paths on top of the main encoder and decoder. This gives the architecture functionality of deep supervision designed to cope with optimization difficulties when training deep networks with limited training data and enable learning of better features for improved discrimination performance of the network (Supplementary Figure 1). In contrast with M-Net which uses a 3D to 2D transformer prior to the M-Net to make use of the volumetric datasets, we adapted the whole network to 3D using 3D convolutional layers.

All networks were implemented in Keras 2.2.4 [27] with TensorflowGPU v1.12.0 backend [28]. We refer readers to original papers for more detailed description of the cited architectures.

Models were trained as to maximize the Weighted Dice Loss which measures the overlap between the true mask and prediction of the model (See Evaluation Metrics). By weighting, we mean focusing the model on the contours by penalizing misclassification at borders of the left ventricle more than the misclassification at the center of the ventricle or in the background. Similarly to [13], the weights at each voxel were calculated based on their distance to the mask border following the equation:

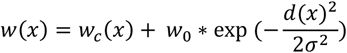

where w_c_ Ω→R is the weight map to balance the class frequencies, d: Ω→R denotes the distance to the ventricular border. We treated σ as a hyperparameter during optimization and set w0 to 2. The weight maps were generated before training for all training image stacks and were supplied to networks together with the original images and their respective masks to supervise the learning process. The output of the models is a mask of the same dimensions as the input image that labels each voxel as either belonging to the left ventricle or the background.

Network hyperparameters have potential to significantly affect the models’ performance. Random search [29] was performed to tune the proposed architectures. Ranges of hyperparameter values used during the tuning were based on previous work and are detailed in Supplementary Table 1. During the hyperparameter optimization, number of down-sampling block and number of convolutional layers within these blocks seemed to have the largest effect on the models’ performance. In the final optimization steps, we fixed the remaining hyperparameters and solely fine-tuned the number of block and layers within the networks. All resulting networks were trained using an SGD optimizer with 0.99 momentum and learning rate of 0.001. σ of 3 was used to calculate the weight maps. We initialized kernels with glorot normal initialization. We also used padded 3D convolutions with patch size of (3,3,3) and ELU non-linear activation layers. We used the batch normalization layers rather than dropout for regularization [30]. The last layer within the networks was followed by sigmoid activation function.

The final segmentation results are obtained using an ensemble of six best-performing models (Figure 2) selected during hyperparameter optimization based on 10-fold cross-validation. To average the predictions of six models, we use a simple majority vote strategy, i.e. at least half of the models must agree on every voxel to consider it a mask. Individual predictions are summed and divided by the number of models. The voxel-wise average prediction above the 0.5 threshold are considered a mask, all other voxels are assigned to the background.

**Fig. 2:**
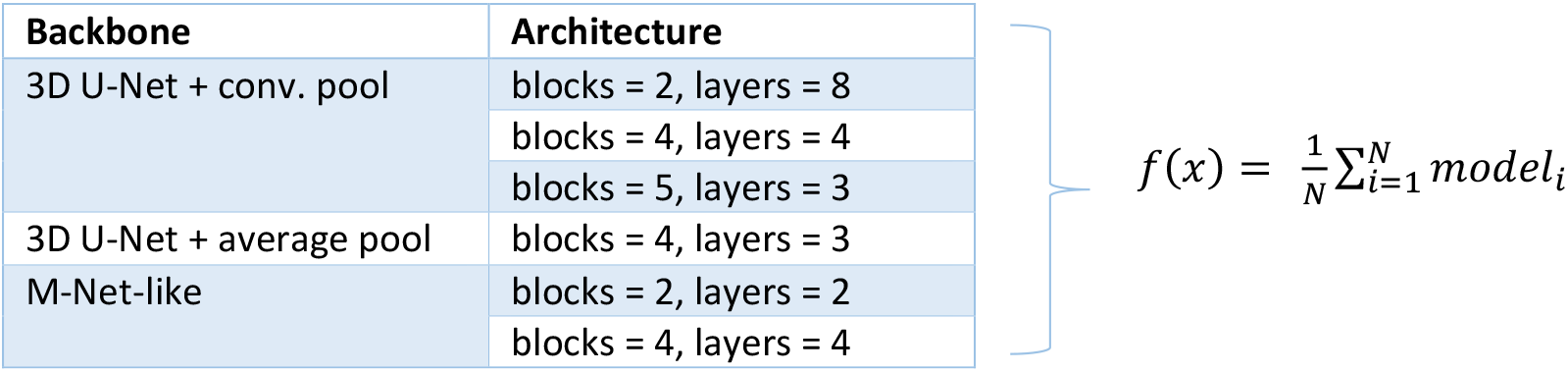
Model ensemble description. Except for the number of down-sampling block and number of layers within the blocks, all other hyperparameters were the same. The two 3D U-nets differ in their pooling strategy used either 3D convolutional layer with stride 2 or 3D average pooling at down-sampling steps. Final segmentations were obtained by a majority vote by independent models.

### Phase Selection

The resulting ensemble performs well on the whole heart cycle instead of end-systole and end-diastole phases exclusively. To select the diastole and systole automatically, we first use predicted segmentations to calculate left ventricular volumes for all phases and create a time-volume curve along the entire heart cycle. We then fit a 4th degree polynomial to the calculated data points. The end-diastole is the phase of largest relaxation and end-systole is defined by the strongest constriction of the ventricle, the peak and the groove points of the fitted curve are considered end-diastole and end-systole volumes, respectively. As the number of recorded phases is finite and phase selection in this manner can select a point which lies between the recorded phases, to match the end-systole and end-diastole selection to a specific scanned 3D image, we select the closest whole phase the one proposed by the curve-fitting approach. The volume of the left ventricle in these two time frames is considered the final end-systole and end-diastole volume.

### Evaluation Metrics

The performance of the ensemble is evaluated from both quantitative and qualitative points of views. Firstly, quality of the resulting segmentations in ED and ES phases was assessed using Dice score [31]. The Dice score is a simple and popular measure of spatial overlap between two segmentation target regions, which can be applied to studies of accuracy in image segmentation and inter-reader variability between different human readers. Dice score is defined as:

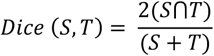

where ⋂ is the intersection between the predicted segmentation S and the true target mask T. Dice score takes values between 0 and 1, with higher values indicating better spatial overlap of the two segmentations.

We also evaluate segmentation accuracy in terms of end-diastole volume (EDV) and end-systole volume (ESV) derived from image segmentations. The LV volumes are calculated by summing up the number of voxels belonging to the corresponding LV mask in the segmentation multiplied by the volume per voxel. Ejection fraction (EF), a critical descriptor of cardiac function, is then calculated using the LV volumes as:

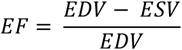

All volumes are expressed in μL and EF is expressed in percentage. The three metrics are calculated for automatic segmentations of the independent testing set. We evaluate mean absolute difference between the calculated parameters and measurements obtained from manual segmentations performed by expert reader.

Additionally, we study the agreement between the automated and human reader measurements using a graphical Bland-Altman analysis [32] and visualize the percent difference between two paired measurements against their average. For instance, in case of EF, the difference is (*EF*_*Reader* 1_ − *EF*_*Automated*_) ∗ 100/*EF*_*Reader* 1_ plotted against the average of each pair of values in the independent testing set. Mean difference between two methods is used to construct limits of agreement which are defined as mean difference ± 1.96 standard deviations of the differences. Such analysis is useful to evaluate the magnitude of the systematic difference, or bias between two methods, to estimate an agreement interval between them, and to uncover possible outliers or potential relationships between the differences and the magnitude of the measurements [33].

To evaluate how well the automatic segmentation compares to the inter-variability between two human readers we performed the same Dice score, EDV, ESV, and EF evaluation and Bland-Altman analysis on manual segmentations performed by two human readers.

## Results

### Dice Score

The automated short-axis LV segmentation was in good agreement with manual segmentation by a human reader at both ED and ES phases. Figure 3 illustrates the exemplary LV segmentation. Compared to the expert segmentations on the pre-selected ED and ES phases, the mean Dice score ± SD for automated segmentation vs expert Reader 1 was 0.93±0.05 (0.92±0.02 for ES and 0.95±0.01 for ED) on the independent testing.

**Fig.3.**
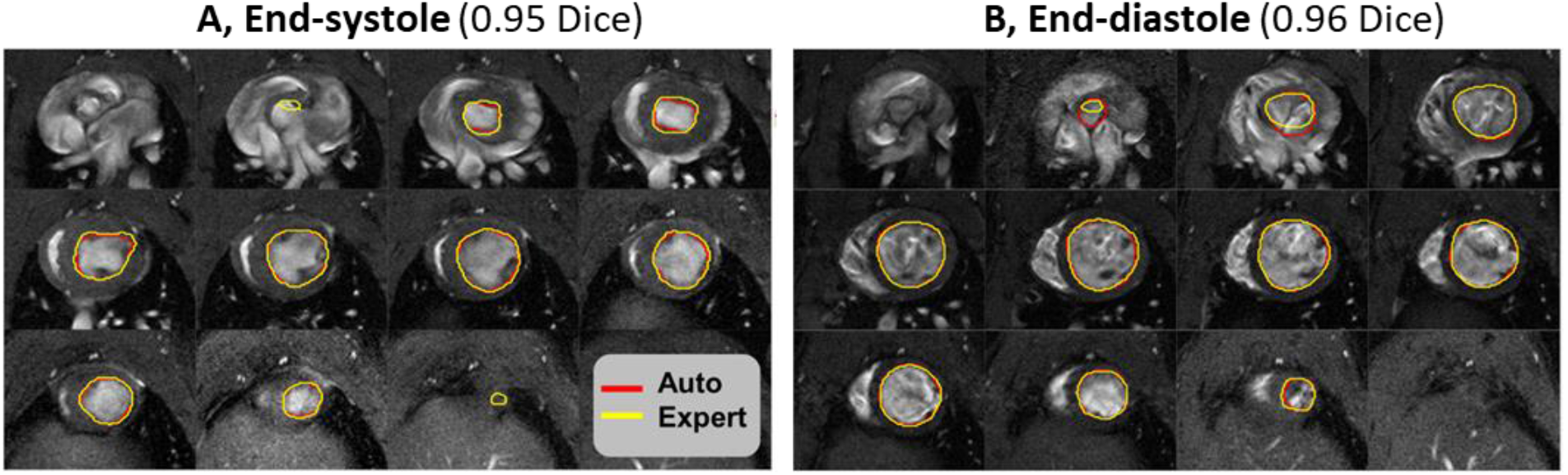
Exemplary segmentation result for short-axis rat MRI images in diastole. Yellow contours show the manual segmentation of the left ventricle. Contours in red show the automatic segmentation. Only end-systole and end-diastole phases are manually annotated. The automated approach is able to segment whole heart cycle.

To evaluate human performance, we also compared the Dice score between manual segmentations by two human readers. Automate vs Reader 1 results were higher than the spatial overlap agreement between two human readers which achieved mean Dice score of 0.90±0.07 (0.87±0.03 for ES and 0.93±0.02 for ED) (Table 1) demonstrating that the automated-human difference is close to or even smaller than the human-human difference in terms of Dice score.

**Table 1:** Dice score evaluation for Automated-Reader 1 vs. Reader 1-Reader 2 segmentations. Automated segmentation agrees well with manual segmentation by expert reader and achieves higher Dice score than the comparison between manual segmentations performed by two human readers.

|  | Automated vs Reader 1 | Reader 1 vs Reader 2 |
| --- | --- | --- |
| Mean Dice | $0.93 \pm 0.05$ | $0.90 \pm 0.07$ |
| ES Dice | $0.92 \pm 0.02$ | $0.87 \pm 0.03$ |
| ED Dice | $0.95 \pm 0.01$ | $0.93 \pm 0.02$ |

Note that cardiac phases other than ES and ED were not considered in the scores as the manual segmentations were not available for the Dice score comparison. Upon visual inspection, 4D segmentation along the time axis yielded convincing results, which were smooth and robust in time. Qualitatively, the 4D segmentations yielded convincing results, which were smooth and robust along the phases. Segmentation of a 3D volume by an ensemble across all recorded phases takes ~ 1.6 min/animal.

### EDV, ESV, and EF evaluation

Next, we evaluate the accuracy of EDV, ESV, and EF measurements directly derived from the automated segmentations. To achieve this, we first need to automatically select the ED and ES phases and determine LV ejection from the calculated volumes.

We use the whole cycle segmentations to calculate the LV volumes at all heart cycle phases. The phase selection is based on fitting a polynomial to the resulting volume curves and is in good agreement with the phase selection performed by human reader (Figure 4). The selection fails for four end-diastole datasets. In the failed cases, the largest volume is assumed the end-diastole in subsequent EDV, ESV, and EF analyses.

**Fig.4.**
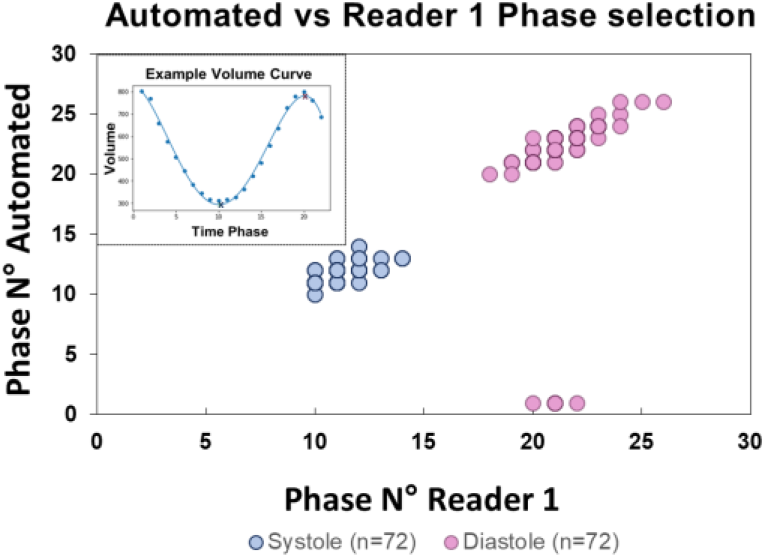
Comparison of Automated vs Reader 1 ED and ES phase selection. Inset figure is an exemplary volume curve from which the ES and ED phases were selected. In case of failed selection, phases with largest and smallest volumes were selected as ED and ES respectively.

Table 2 reports mean absolute difference between automated and manual measurements and between measurements by two expert readers in terms of EDV, EDV, and EF mean relative difference evaluated in the testing set. The automated segmentation leads to a higher accuracy when comparing automated-reader vs. reader-reader with the mean relative difference of 5.0±4.4 % vs. 8.2±4.6 % for EDV, 8.2±6.1% vs. 25.9±9.1% for ESV, and 6.3±2.4% vs. 6.8±2.3% for EF.

**Table 2:** Mean absolute differences for Reader 1-Automated vs. Reader 1-Reader 2. Computer-human difference is comparable to the inter-reader variability for the derived metrics.

|  | Reader 1 vs Automated | Reader 1 vs Reader 2 |
| --- | --- | --- |
| $\Delta$ EDV (μL) | $43.4 \pm 27.4$ | $59.3 \pm 31.8$ |
| $\Delta$ ESV (μL) | $35.6 \pm 22.3$ | $79.4 \pm 25.8$ |
| $\Delta$ EF (%) | $6.2 \pm 2.3$ | $6.8 \pm 2.4$ |

### Bland-Altman analysis

To graphically compare agreements between automated-human and human-human, we visualize EDV, ESV, and EF mean percent differences in a Bland-Altman plot (Figure 5). The plots indicate systematic difference between the methods in both automated-reader and reader-reader comparisons. The relative bias of the automated method vs human reader is similar to that between two humans in case of EF but is lower for EDV and ESV. We observe weak relationship between the differences and the magnitude of EF for automated vs human. Overall, our automated method performs on par with human reader and leads to slightly better agreement with manual segmentation compared to the agreement between two human readers.

**Fig.5:**
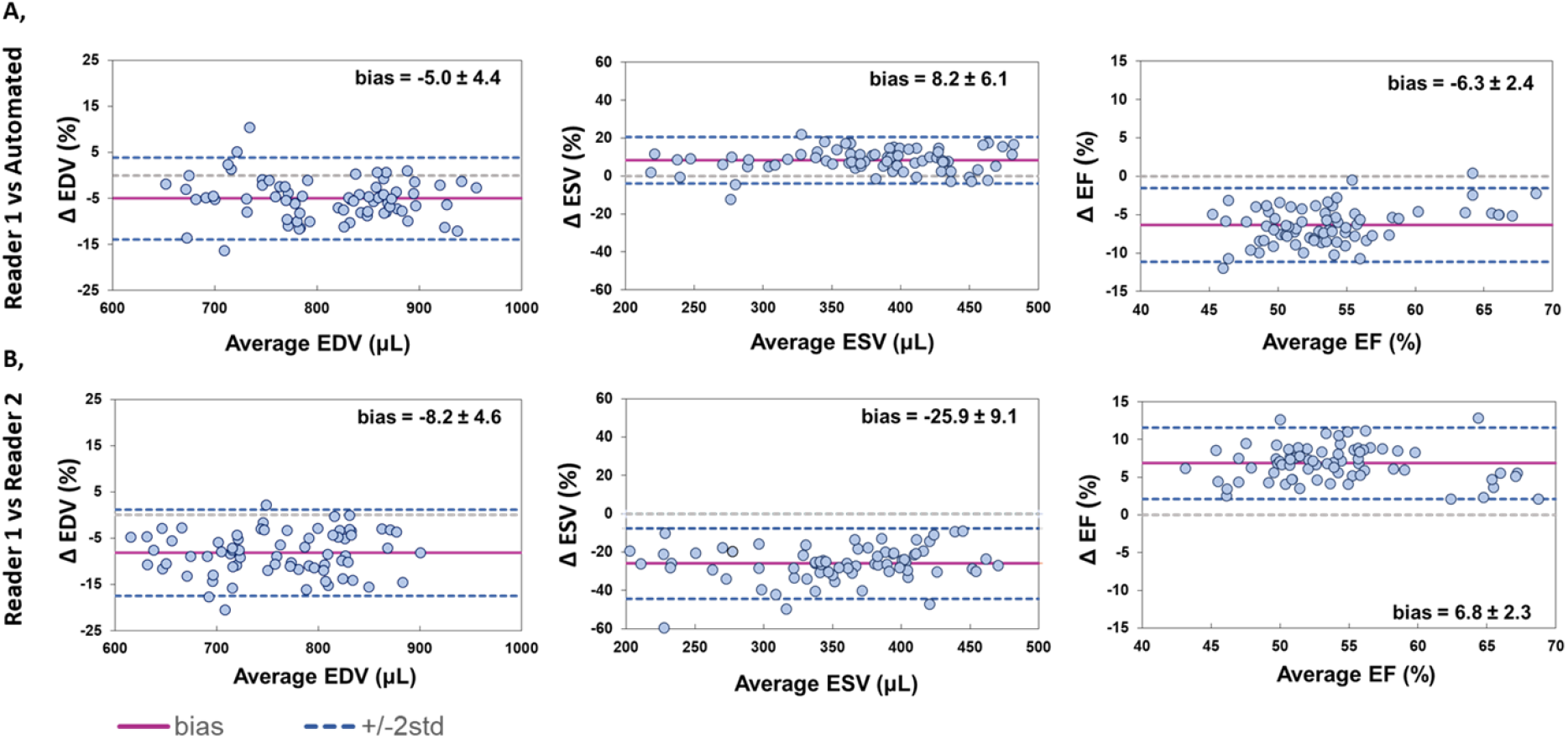
Bland-Altman plots for Reader 1 vs. automated method (A) and Reader 1 vs. Reader 2 (B) for end-diastole (right), end-systole (middle), and ejection fraction (left). The figures show the agreement between the methods on a testing set of 72 subject by plotting the differences between two methods against the average of the corresponding measurements. The purple horizonal line represents the mean percent difference, or bias; blue dotted lines are limits of agreement denoting ±1.96 standard deviations from the mean difference, gray line is the theoretical line of equality for the two methods.

Overall, we present an efficient ensemble method for end-to-end segmentation of short-axis LV volumes from rodent CMRI datasets on par with human experts in terms of quality of segmentations assessed with Dice score and accuracy of derived left ventricular volumes and ejection fraction. More extensive validation is required, but the method has a great potential for improving work efficiency of analysis teams in the future by creating automated reports of parameters of interest for animals in the study.

## Discussion

Due to the time constraints imposed by the manual segmentation process, current routine analysis requires only segmentations at two phases of the cardiac cycle, end-systole and end-diastole. In the future, robust LV delineation of all phases of the heart cycle could allow more detailed LV assessment in temporal volumes Our tool seems to perform satisfactorily in this respect and presents a first step towards full cycle analysis of rodent heart function.

Regarding the segmentation quality, most errors appear at the apical slices of the LV region. This occurs mostly in end-systole, where in rare cases, the ensemble fails to detect the presence of LV completely, resulting in a lower mean Dice score compared to end-diastole. This observation is consistent with results in other works and is likely due to the higher complexity and variability of the LV appearance in end-systole compared to end-diastole [23]. Additionally, small LV size on most apical and basal slices leads to high imbalance between number of voxels that belong to the LV region compared to the background which could affect the segmentation quality in these regions. Moreover, as the left ventricle is generally smaller in volume, smaller misclassifications can translate to disproportionally lower Dice scores than in the end-diastole. Some of the other segmentation errors such as added regions or inside and boundary holes described in the past were also observed. We believe these can be resolved by finer tuning of the model ensemble. Currently, our ensemble method uses model averaging so that each ensemble member contributed equally to final segmentations. In the future, we would like to explore the use of larger model ensemble or the weighing of the ensemble that would allow the contribution of each model to be weighted proportionally to the trust we assign to further improve the segmentations.

The ensemble performed on par with human expert in terms of agreement between EDV, ESV, and EF derived from automated vs manual segmentations and is similar to inter-reader variability between two human readers with the mean relative difference of 5.0±4.4 % vs. 8.2±4.6 % for EDV, 8.2±6.1% vs. 25.9±9.1% for ESV, and 6.3±2.4% vs. 6.8±2.3% for EF. However, bias between the methods seems to be a problem in both automated-human and human-human cases. In case of consistent bias, some authors recommend adjusting the results post-hoc by subtracting the mean difference from the new method for the final EDV, ESV, and EF reporting. Currently, our method is trained with annotations from 2 different readers, which could explain slightly lower systemic bias in case of automated-human analysis in all three parameters. To improve the segmentation bias further, we will aim to include annotations from multiple additional readers. This could potentially allow the ensemble to learn a better consensus estimate across the group of observers and become susceptible to biases.

Another limitation of our work linked to the problem of insufficient variability of our training dataset is that the ensemble was trained on images acquired using the same standard imaging protocol and scanner model. This suggests that our method might lack the generalization capabilities when presented with data from different scanner, acquired using different imaging protocol, or including more abnormal previously unseen LV pathologies. The ensemble might need to be fine-tuned when new, sufficiently different data is acquired which will require manual labelling of some of the new data to improve the segmentations. This limitation might prevent the model from being successfully deployed in the analysis pipeline and therefore diminish its impact for improving the preclinical workflow.

In the future, it would be useful to explore whether we could create a sufficiently heterogenous multi-scanner dataset for training and evaluation of automated models that could cover typical rodent CMR imaging protocols and LV pathologies in a fashion similar to large scare efforts in human CMR imaging.

However, this approach may not scale well, as it requires the organized collection and labeling of a large dataset covering variety of possible cases. Rather than collecting and segmenting new datasets manually from scratch, the existing ensemble could be used to pre-segment the images with human experts adjusting the segmentations before adding them to the training set. Other approaches such as weakly and semi-supervised learning, self-supervised learning and unsupervised learning might be more realistic [10]. Several studies have focused on adopting more complex data normalization and data augmentation strategies to simulate various data distributions to improve robustness of the CMRI segmentation models and enable better generalizability to new data. We would like to explore further affine and elastic data augmentation strategies, random noise addition, and image contrast adjustment in the future to create a training set that could better reflect the spectrum of the real data distribution and create more robust generalizable methods for segmentation of wide ranges of rodent CMR images.

## Conclusion

We present a successful application of deep learning to automated end-to-end segmentation of rodent left ventricle from CRMI datasets. Quantitative and qualitative evaluations and inter-reader variability assessments demonstrate human-level performance of our method on the independent testing set. Overall, our implementation allows automated extraction of clinically relevant parameters with minimal input of human users that has potential to translate into substantial saving of time and resource and could significantly speed up reporting of results to analysis teams. Furthermore, the present work may also find its relevance in other applications, for example for the segmentation of mice rather than rat datasets. We anticipate this to be a starting point for automated CMR analysis facilitated by deep learning in our team. In the future, AI will be essential when assessing pharmacological interventions designed to improve cardiac function with increased speed and high precision.

## Acknowledgements

The authors acknowledge that the work was fully sponsored by AstraZeneca, Innovative Medicines and Early Development, Gothenburg, UK.

## Supplementary Materials

**Supplementary Table 1:** Hyperparameter ranges for hyperparameter optimization. *Sigma weighting parameter when weighted dice loss was used for training.

| Hyperparameter | Range of values |
| --- | --- |
| Activation Function | Elu, Relu, Prelu |
| Optimizer | Adam, RMSprop, SGD (with momentum=0.99) |
| Loss | Binary cross-entropy, Dice Loss, Weighted Dice Loss |
| Filter Size | (3,3,3), (5,5,5), (10,10,10) |
| N° of Filters | 8, 16, 32, 64 |
| Batch Normalization | True, False |
| Dropout | 0.0, 0.5, 0.9 |
| N° of down-sampling blocks | 1, 2, 3, 4, 5 |
| N° of layers per block | 1, 2, 3, 4, 5, 8, 10 |
| Decay | 0.0, 0.25, 0.5 |
| Batch Size | 1, 2, 5, 10 |
| Learning Rate | 0.001, 0.01, 0.1 |
| Sigma* | 0.5, 1, 2, 3, 4, 5, 7, 10 |

**Supplementary Fig. 1:**
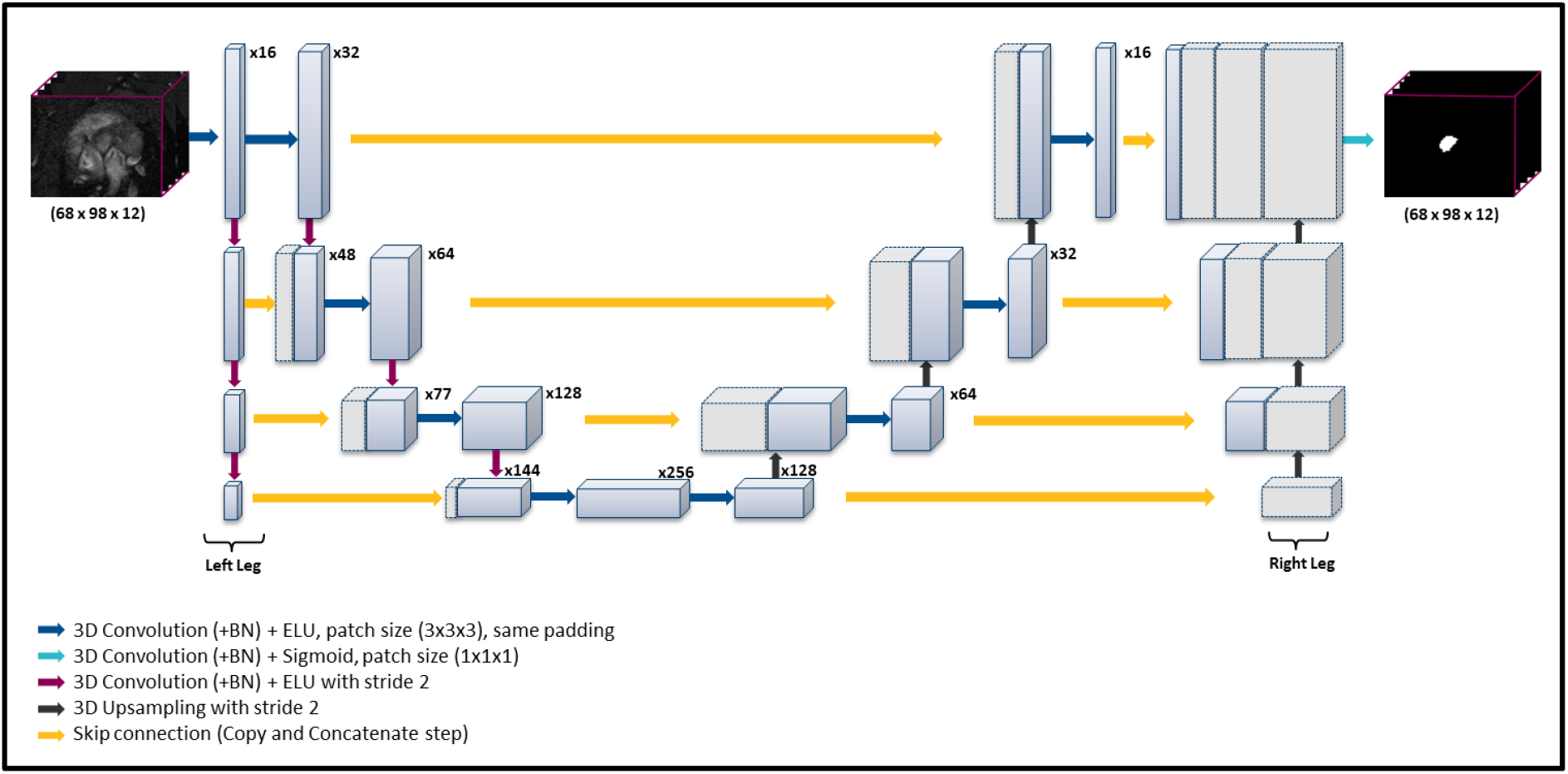
Schematic representation of the M-net CNN architecture. Blue boxes represent feature maps with number of maps denoted near the box. Gray boxes represent copied feature maps. For simplicity, we have reduced the number of down-sampling and up-sampling blocks as well as number of layers per block in the diagram.

